# Dose-dependent effects of inhaled corticosteroids on bone mineral density in postmenopausal women with asthma or COPD: A registry-based cohort study

**DOI:** 10.1101/184937

**Authors:** Wenjia Chen, Kate M. Johnson, J. Mark FitzGerald, Mohsen Sadatsafavi, William D. Leslie

## Abstract

**Background:** The effect of long-term inhaled corticosteroid (ICS) therapy on the bone health of older adults remains unclear due to its possible impact on bone mineral density (BMD).

**Objective:** To evaluate, cross-sectionally and longitudinally, the impact of ICS use on BMD in postmenopausal women with asthma or chronic obstructive pulmonary disease (COPD).

**Methods:** We used a population-based bone densitometry registry linked with administrative health data of the province of Manitoba, Canada (1999–2013), to identify women with diagnosed asthma or COPD. ICS use was defined as cumulative dispensed days prior to baseline BMD (cross-sectional analysis), and medication possession ratio (MPR) between two BMD measurements (longitudinal analysis). Results were adjusted for multiple covariates including the underlying respiratory diagnosis and its severity.

**Results:** In the cross sectional analysis, compared with non-users, women with the highest tertile of prior ICS exposure had lower baseline BMD at the femoral neck (-0.09 standard deviations [SD] below a healthy young adult, 95% CI: −0.16, −0.02) and total hip (-0.14 SD, 95% CI: −0.22, −0.05), but not at the lumbar spine. Longitudinally, the highest tertile of ICS exposure was associated with a slight decline in total hip BMD relative to non-users (-0.02 SD/year, 95% CI: −0.04, −0.01), with no significant effect at the femoral neck and lumbar spine. Middle and lower tertiles of ICS use had no significant effects.

**Conclusion:** High exposure to ICS was associated with a small adverse effect on baseline hip BMD and total hip BMD loss in post-menopausal women with asthma or COPD.

---

**What is the key question?** What is the safety of long-term use of inhaled corticosteroids in postmenopausal women with chronic respiratory disease?

**What is the bottom line?** Postmenopausal women with over 50% adherence to inhaled corticosteroids tend to have slightly accelerated bone mineral density loss at the total hip, but overall this loss was very minor.

**Why read on?** For clinicians making treatment decisions that must balance efficacy and risk of side effects, this study provides a population-based assessment of the long-term dose-response association between inhaled corticosteroids and bone mineral density, and highlights the need to maintain minimally effective doses in this patient group.

## INTRODUCTION

Inhaled corticosteroids (ICS) are commonly used in the management of chronic diseases of the airways due to their impact on airway inflammation. ICS reduce the rate of exacerbations, decrease respiratory symptoms, improve lung function and quality of life in patients with asthma^1^. The use of ICS in chronic obstructive pulmonary disease (COPD) is less well established, but it has been shown to reduce exacerbations in moderate to severe COPD, especially in combination with a long acting beta agonist^2,3^.

Despite its efficacy, the safety of long-term ICS use remains contentious. In particular, ICS use is strongly associated with an increased risk of pneumonia^4^, and more moderately associated with decreases in bone mineral density (BMD)^5-9^, and an increased risk of fractures^10^, although several meta-analyses have found no effect^4,11^. The relationship between ICS and BMD is generally found to be dose-dependent^5,6,9^, however, the typical dosage assessed in these studies is high. A population-based assessment of long-term ICS use at a wide range of doses would be relevant from both a pathophysiological perspective and to clinicians making treatment decisions that must balance safety and efficacy. In particular, the dose at which ICS has an impact on BMD might depend on the age and sex of the patient^4,9,12,13^, and the safe dose may be lower in populations in which natural bone loss is more pronounced, such as in postmenopausal women^14^. However, the association between ICS use and BMD decline has not been well studied in this population. There is evidence that the impact of ICS on BMD is greater in postmenopausal than in premenopausal women^8^, although the sample sizes of studies of postmenopausal women have tended to be small^7,8^. More detailed evaluation of the dose-response relationship between ICS therapy and BMD in postmenopausal women with chronic airway diseases is needed to determine whether preventative therapy is necessary to reduce BMD decline and the risk of fractures.

The objective of this study was to examine the impact of ICS on BMD loss in postmenopausal women with asthma or COPD in routine clinical practice. We hypothesized that in women with asthma or COPD, after controlling for disease severity and patient characteristics, BMD is lower in those exposed to ICS compared with unexposed women, and that BMD declines more rapidly with increasing exposure to ICS.

## METHODS

### Data sources

The province of Manitoba, Canada, provides universal health care to its population of 1.3 million residents (as of 2016) ^15^. The administrative needs of maintaining the public health care system have resulted in the creation of centralized health care databases, which comprehensively capture information about hospital discharges, physician billing claims, prescription medication dispensations as well as demographics, registration and vital statistics. These databases have low rates of missing data and high validity^16,18^. The current study was based on bone densitometry services provided between April 1, 1999 and March 31, 2013 under a province-wide bone densitometry program^19^. The population-based clinical BMD registry records information related to all bone densitometry services in the province since 1990 (completeness and accuracy≥99°%)^20^. The BMD registry was linked at the individual level with other population-based provincial health administrative data held by the Manitoba Centre for Health Policy Data Repository via an encrypted personal health number. The study was approved by the Human Research Ethics Board of the University of Manitoba. Data access permission was obtained from the Manitoba Health Information Privacy Committee.

### Study population

This retrospective cohort study had both cross-sectional and longitudinal components. Figure 1 displays the schematic presentation of the study design. The study population consisted of women who were at least 40 years of age, had continuous health care coverage for at least 3 years prior to undergoing their first BMD test, and had a previous diagnosis of asthma or COPD. These diagnoses were identified by the presence of one or more hospitalizations or two or more physician claims with diagnostic codes for asthma or COPD, during the 3-year period prior to the first BMD test. Asthma-specific inpatient and outpatient encounters were determined based on International Classification of Diseases, 9^th^ Edition (ICD-9) codes of 493.x, and ICD-10 codes of J45.x, J46.x. COPD-specific encounters were determined by ICD-9 codes of 491.xx, 492.xx, 493.2x, 496.xx, and ICD-10 codes of J43.xx, J44.xx. For each patient, the *index date* was defined as the date of first (baseline) BMD measurement.

**Figure 1.**
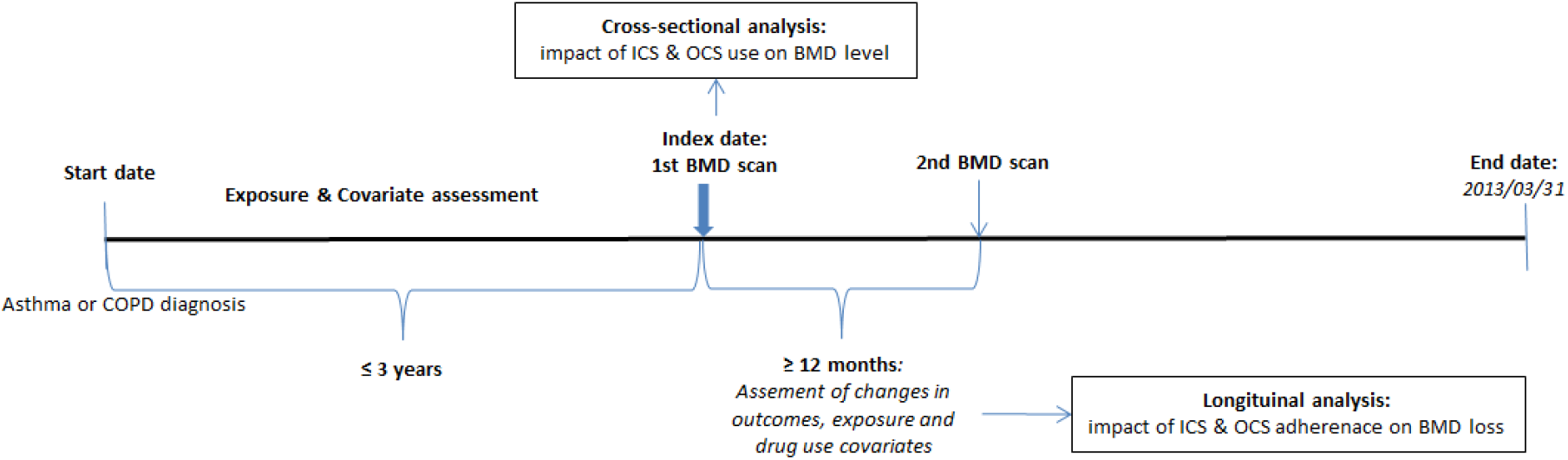
Schematic presentation of study design. BMD, bone mineral density; COPD, chronic obstructive pulmonary disease, ICS, inhaled corticosteroids; OCS: oral corticosteroids.

### Outcomes

BMD testing was performed using dual-energy X-ray absorptiometry scans of the hip and spine with a pencil-beam instrument (Lunar DPX; GE Lunar, Madison WI, USA) prior to 2000 and fan-beam instruments (Lunar Prodigy or iDXA; GE Lunar) afterwards. The program’s quality assurance is under strict supervision by a medical physicist^19^. Instruments were cross-calibrated and no clinically significant differences were detected^20^ The instruments used for this study exhibited stable long-term performance (coefficient of variation <0.5%). All reporting physicians and supervising technologists are required to maintain DXA certification with the International Society for Clinical Densitometry (ISCD).

BMD was measured at the femoral neck, total hip, and lumbar spine (L1-4). The primary site for the cross-sectional analysis was the femoral neck as this is the reference standard for description of osteoporosis diagnosis and for fracture risk assessment^21^. To examine the cross-sectional association between ICS use and baseline BMD, we reported BMD as measured in the first scan in T-scores (i.e., the number of standard deviations above or below the mean of a healthy young adult white female^22^). Hip T-scores were calculated using U.S. National Health and Nutrition Examination Survey (NHANES) III reference values^21^. Lumbar spine T-score calculations used manufacturer’s reference data for U.S. white females^23^.

We examined the longitudinal loss in BMD between the first and second scans as the change in T-scores divided by the time in years between the two scans where the second scan was at least 12 months after the baseline examination. The primary site for the longitudinal analysis was the total hip since it has the best test-retest precision and is the least affected by age-related degenerative artifact^24^.

### Exposures

All exposure measures were obtained from the comprehensive provincial pharmacy system using data from the Drug Program Information Network (DPIN)^17^. The use of ICS was measured in multiple ways. Cumulative dispensed days (primary exposure) and total dispensed quantity (mcg of beclomethasone equivalent, secondary exposure) of ICS use prior to the index date were used for the cross-sectional analysis. For the longitudinal analyses, the dispensed days between the two BMD measurements, measured by medication possession ratio (MPR) was used. MPR was defined as the ratio of days that a patient was on medication divided by the total number days observed for that patient (a value between 0 and 1). As such, its definition is independent of the length of the time window. As a secondary longitudinal exposure, we also measured total dispensed quantity of ICS between the two scans, which was normalized for time (divided by the time interval between the two scans).

For each exposure definition, based on the tertiles of exposure, ICS use was classified into four categories: none, lowest-, middle- and highest tertile. The reference category was none (no use).

### Statistical analyses

All analyses were performed with Dell Statistica (Version 13.0, Dell Inc. 2015). A 2-sided P-value of 0.05 was set as the threshold for assessing statistical significance.

For the cross-sectional analysis, we used generalized linear models with analysis of covariance to estimate the association between the history of ICS use and the levels of BMD at baseline, with parallel analyses performed for each of the three different BMD measurement sites. We used BMD T-score as the outcome and tertiles of ICS use prior to the first BMD scan as the main exposure. We adjusted for major respiratory diagnosis (COPD or asthma), covariates from the Fracture Risk Assessment Tool (FRAX^25^): age, body mass index (BMI), self-reported parental hip fracture and smoking status on the index date, as well as history of major fracture, rheumatoid arthritis (based on ICD codes), and high alcohol intake (alcohol/substance abuse diagnosis, based on ICD codes) assessed prior to the index date from administrative data. We also adjusted for use of osteoporosis medications (bisphosphonates, calcitonin, systemic estrogen products, raloxifene, teriparatide). In addition, to account for the potential confounding effect of disease severity, we also adjusted for total number of dispensed days (in tertiles) of oral corticosteroids (OCS, expressed as the MPR between the two tests), and the number of asthma/COPD-related hospitalizations and physician visits in the 3 years prior to the index date. We tested the interaction effects of disease diagnosis and ICS use on BMD loss in an exploratory analysis. In a sensitivity analysis we replaced days of ICS use with total dispensed quantity of ICS (beclamethasone-equivalents) as the alternative exposure.

For the longitudinal analysis, we regressed the effects of ICS (primary exposure: MPR, secondary exposure: time-adjusted total dispensed quantity) on the longitudinal annual changes in BMD (expressed as T-score SD/year) between the two consecutive BMD tests, with parallel analyses performed for changes in all three BMD measurement sites. We conducted a sensitivity analysis in which we repeated the longitudinal analysis for the subgroup of women who did not have any estrogen or osteoporosis medication exposure during the observation period.

## RESULTS

### Cross-sectional analysis: association between history of ICS use and baseline BMD

The cross-sectional sample included 6,561 female patients aged 40 years and above, including 63% with a primary diagnosis of COPD and 37% with asthma, respectively (Table 1). The average age at baseline was 65.2 years (SD=10.8). Approximately, 51% of patients had ever used ICS prior to BMD testing and were divided into 3 tertiles based on total days of usage (lowest tertile: 1-155 days of use, middle: 156-719 days, highest: above 720 days). The mean T-scores of hip, femoral neck and lumbar spine were −1.0, −1.5 and −1.1, respectively. Based on the lowest score across the sites, osteoporosis was prevalent in 31% of patients at baseline.

**Table 1.** Descriptive characteristics of the study sample

|  | Overall |
| --- | --- |
| **Cross-sectional sample (n=6,561)** |  |
| Diagnosis, n% |  |
| *COPD* | 4,110 (62.6) |
| *Asthma* | 2,451 (37.4) |
| Age, years | 65.2 ± 10.8 |
| Body mass index, kg/m<sup>2</sup> | 28 ± 6.3 |
| Prior fracture, n% | 1,119 (17.1) |
| Rheumatoid arthritis | 251 (3.8) |
| High alcohol intake | 380 (5.8) |
| Current smoker | 936 (21.1) |
| Parental hip fracture | 565 (12.7) |
| Any ICS use | 2,626 (40.0) |
| Any oral corticosteroid use | 3,274 (49.9) |
| Any osteoporosis drug use | 3,427 (52.2) |
| Lumbar spine T-score | -1.1 ± 1.5 |
| Femoral neck T-score | -1.5 ± 1.0 |
| Total hip T-score | -1.0 ± 1.3 |
| Minimum site osteoporotic<sup>†</sup> | 2,050 (31.2) |
| Days of prior ICS use, n% |  |
| None | 2,626 (40.0) |
| Lowest tertile (1-155 days) | 1,323 (20.2) |
| Middle tertile (156-719 days) | 1,301 (19.8) |
| Highest tertile (>719 days) | 1,311 (20.0) |
| Days of prior OCS use, n% |  |
| None | 3,274 (49.9) |
| Lowest tertile (1-90 days) | 1,149 (17.5) |
| Middle tertile (91-365 days) | 1,052 (16) |
| Highest tertile (>366 days) | 1,086 (16.6) |

| **Longitudinal sub-sample (N=1807)** |  |
| --- | --- |
| Diagnosis, n% |  |
| *COPD* | 1,066 (59.0) |
| *Asthma* | 741 (41.0) |
| BMD interval, years | 4.8 ± 2.4 |
| Change in lumbar spine T-score, SD/y | 0.011 ± 0.156 |
| Change in femoral neck T-score, SD/y | -0.027 ± 0.098 |
| Change in total hip T-score, SD/y | -0.025 ± 0.112 |
| Adherence to ICS, MPR, n% |  |
| None | 890 (49.3) |
| Lowest tertile (≤0.16) | 302 (16.7) |
| Middle tertile (0.17-0.50) | 303 (16.8) |
| Highest tertile (>0.50)) | 301 (16.7) |

Values are mean ± standard deviation or n (%).
COPD, chronic obstructive pulmonary disease.
\*P-values were obtained from student T-test.

Values are mean ± standard deviation or n (%).

COPD, chronic obstructive pulmonary disease.

^*^P-values were obtained from student T-test.

^†^Osteoporosis was defined as −2.5 or lower in T-scores based on the minimum T-score obtained from the three sites.

Figure 2 shows the association between total days of ICS use prior to BMD testing and the BMD T-scores at baseline. Prior ICS exposure was associated with lowered baseline T-scores for the femoral neck (p=0.005) and total hip (p=0.002), but not for the lumbar spine (p=0.12). Specifically, the highest tertile of prior ICS days was associated with lowered T-scores compared to no ICS use in femoral neck and total hip (-0.09 [95% CI: −0.16, −0.02, p=0.009], −0.14 [95% CI: −0.22, −0.05, p=0.001], respectively). The effects of lowest and middle tertiles of ICS use were not significantly different from non-users across three sites. The effects of prior ICS exposure on baseline T-scores did not significantly differ between the COPD and asthma patients (p=0.52 for the interaction term between disease and ICS use).

**Figure 2.**
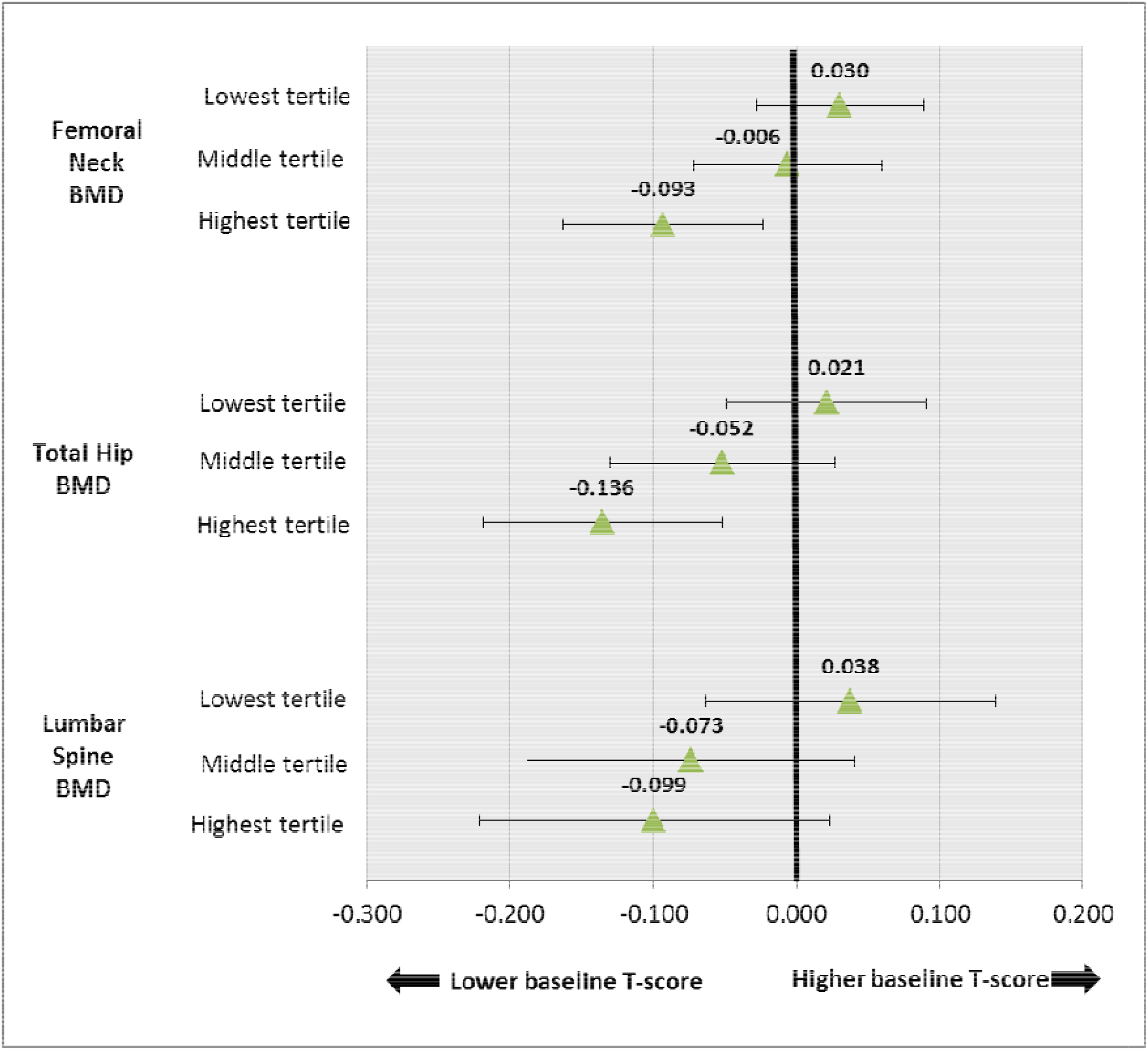
Cross-sectional association between history of ICS use and baseline BMD T-scores from the multiple linear regression models for the (top) femoral neck, (middle) total hip, and (bottom) lumbar spine. Drug use was measured as the total number of dispensed days of ICS before 1^st^ BMD scan and was categorized into tertiles, with the reference group being “No Use”. Lowest tertile, 1-155 days of ICS use, middle tertile, 156-719 days, highest tertile, above 720 days. Error bars show the 95% confidence interval. BMD, bone mineral density; ICS, inhaled corticosteroid.

In the secondary analysis, we changed the main exposures from the total days to the total dispensed quantity of prior ICS use. Results were consistent for all three sites: compared to no use, the highest tertile of ICS quantities (>840,000mcg of beclomethasone equivalent) was associated with lower baseline T-scores for the femoral neck (-0.09 [95% CI: −0.16, −0.02, p=0.006]) and total hip (-0.15 [95% CI: −0.23, −0.06, p<0.001]), but not for lumbar spine (p=0.11).

### Longitudinal analysis: effects of ICS exposure on BMD change

From the initial sample, we identified 1,807 women (59% COPD, 41% asthma) who received a second BMD scan that occurred at least 12 months after baseline scan. The average time interval between the first and second scans was 4.8 years (SD=2.4). ICS were used in 51% of patients between the two scans, with each tertile of ICS MPR comprised of 17% of patients (lowest: <0.16, middle: 0.16-0.50, highest: >0.50). From baseline to the second BMD scan, total hip and femoral neck T-scores had decreased (-0.025 SD/year and −0.027 SD/year, respectively), but the lumbar spine T-score increased (+0.011 SD/year) (Table 1).

Figure 3 shows the longitudinal change in BMD T-scores across levels of ICS use. Overall, ICS use only had a significant effect on the longitudinal decline of BMD at the total hip site (p=0.025) but not at lumbar spine or femur neck (p-values: 0.25, 0.68, respectively). The highest tertile of ICS MPR versus no use was associated with a significant decline in total hip T-score (-0.024 SD/year [95% CI: −0.040, −0.008], p=0.003), whereas the lowest and middle tertiles of MPR had no significant effects. The highest tertile of MPR also led to a borderline decline in lumbar spine T-score (-0.024 SD/year [95% CI: −0.047, 0.000], p=0.050). When women with estrogen or osteoporosis medication exposure were excluded from the sample, the highest tertile of ICS MPR still predicted a borderline loss in total hip BMD T-scores but not in other sites (0.026 SD/year [95%CI: −0.051, 0.000], p=0.047).

**Figure 3.**
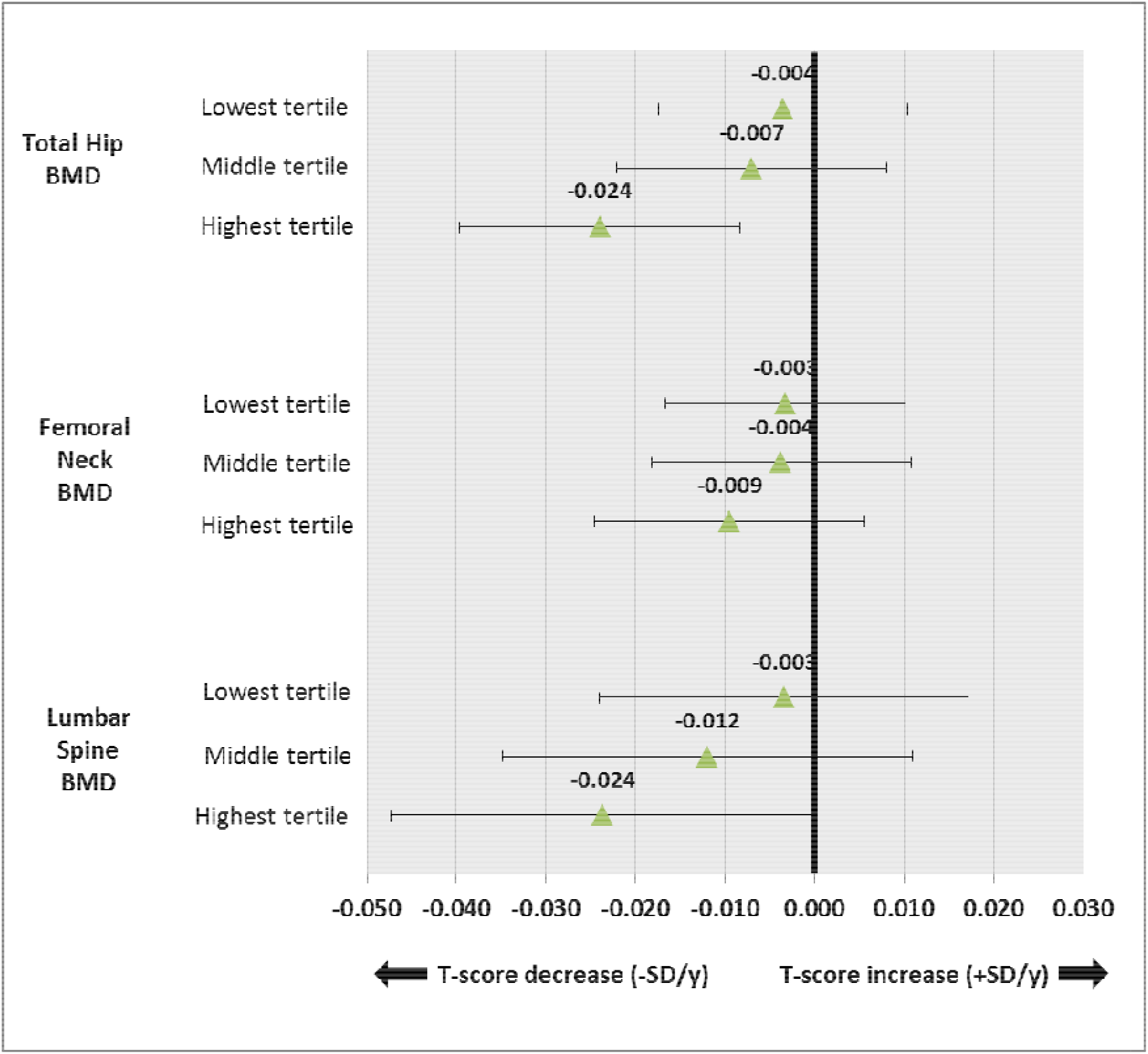
Longitudinal effects of medication possession ratio (MPR) of ICS use on annualized changes in BMD T-scores from the multiple linear regression analysis for the (top) total hip, (middle) femoral neck, and (bottom) lumbar spine. MPR of drug use was measured between 1^st^ and 2^nd^ BMD scan and categorized into tertiles, with the reference group being “No Use”. Lowest tertile: <0.16, middle tertile: 0.16-0.50, highest tertile: >0.50. Error bars show the 95% confidence interval. BMD, bone mineral density; ICS, inhaled corticosteroid, ±SD/y, changes in T-score standard deviation per year.

In the secondary analysis which used time-normalized, between-scan total dispensed quantity of ICS as the exposure variable, results were consistent with the primary analysis: only the highest tertile of ICS quantities (>124,875 mcg of beclomethasone equivalent, time-normalized) was associated with BMD decline in total hip compared to no use (-0.020 SD/year [95% CI: −0.035,0. 004], p=0.016), whereas ICS quantity had no significant effect on other sites.

## DISCUSSION

We used administrative health data of a well-defined geographic area with complete health care coverage linked with a bone densitometry database to examine the cross-sectional and longitudinal impact of ICS use on BMD in postmenopausal women with previously diagnosed asthma or COPD. In the cross-sectional analysis, the highest tertile of ICS use (previous use of more than 720 days) was found to negatively impact BMD at the total hip and femoral neck after taking into account an index of disease severity and common fracture risk factors. In the longitudinal analyses, receiving ICS for more than 50% of the time between the two scans was associated with a modest decline in total hip bone density. Overall, these associations are considered weak. In the cross-sectional analysis, only patients with two or more years of prior exposure to ICS had baseline bone density lower than the non-users, with the T-score reduction much less than one standard deviation for the femoral neck (SD=0.14) and total hip (SD=0.13). Moreover, in the longitudinal analysis, the minor BMD loss (-0.024 SD/year) in total hip among patients with over 50% adherence to ICS (highest tertile) would need to be sustained for over 40 years to produce one standard deviation reduction in total hip BMD.

Our findings are in line with other studies that have found an association between ICS therapy and minor BMD loss^5-9^, and this response is generally observed to be dose-dependent^5,6,9^. In general, the patients in our study were receiving low dose ICS therapy; 85% of patients were dispensed less than 5 puffs (100mcg/puff) of beclomethasone-equivalent per day, although actual medication intake is likely even lower^28^. Similar to Wong et al.^6^, we observed a dose-dependent response between cumulative dispensed quantity and baseline BMD in the cross-sectional analysis, and only the highest tertile of MPR was associated with a decline in BMD. However, the ICS doses observed here are lower than doses that have previously been observed to have a negative effect on BMD^8,9^ or the risk of fractures^29^. Indeed, a pooled analysis of six observational studies found that ICS had a minimal impact on the risk of fractures when the dose of beclomethasone equivalent was below 500 mcg per day^10^. However, women were underrepresented in this analysis, and our study provides some evidence to suggest that the safe dosage may be lower for postmenopausal women than in other populations. Pooled analyses that found no impact of ICS on BMD^11,12^ may have benefited from subgroup analyses within this population at risk of osteoporosis.

The impact of ICS use on BMD varied between bone sites. The hip was the only site at which we observed an effect in both the cross-sectional and longitudinal analyses, and BMD at the lumbar spine was not significantly affected by ICS use in both analyses. These differences may be due to age-related degenerative changes, which are particularly common in the lumbar spine. Although we did not observe a strong association between ICS use and BMD, it is possible that the positive impact of ICS therapy on patient mobility and respiratory function offset its negative impact on BMD and resulted in a smaller net effect. For example, van Staa et al.^30^ found that ICS use increased the risk of fractures in respiratory patients, but respiratory patients not on ICS therapy still had a higher risk of fractures than healthy controls, suggesting the increased risk of fractures was due to the respiratory disease itself rather than ICS use.

Our study has several strengths. First, we assessed ICS use and initial bone density in 6,561 patients, and the change in bone density over an average of five years for 1,087 patients, which is a very robust sample size compared to previous studies^5,8,31^. The registry-based nature of the study sample reduces many issues associated with sample representativeness that are common in cohort studies, including low participation rates, self-selection, and participants lost to follow-up. It is also likely to be very representative of patients in routine clinical practice who are felt to be at increased risk for osteoporosis. In addition, ICS was objectively measured using a prescription drug database, which eliminates bias due to self-reporting. Our sample likely included patients with a wide range of risk factors for osteoporosis. To the best of our knowledge, our study is the first to apply a longitudinal design to a registry-based sample to assess the association between ICS use and BMD. Further, we determined the impact of ICS independent of well-established fracture risk factors, as well as other important predictors of bone density including smoking history and use of osteoporosis drugs or oral corticosteroids. In addition, unobserved, time-fixed confounding effects were accounted for in the longitudinal analysis because BMD comparisons were made within patients. This helped enable inference on the causal effects of ICS use on progressive BMD loss.

However, our study also has several limitations. First, we were unable to perform rigorous adjustment for lung function or the level of systemic inflammation as potentially important confounders because these parameters were unavailable in the data. These factors can change rapidly over time and might independently affect BMD. However, we did adjust for disease severity based on the intensity of resource use for respiratory conditions, which might account for part of the longitudinal variation in lung function and inflammation. Second, our sample consisted of patients for whom a BMD scan was requested by their physician. As a result, the majority of patients with COPD were osteopenic, and the average age of asthma patients was older (65 years) than is typically observed asthma cohorts. Our findings may not apply to younger postmenopausal women, or patients who are not already at risk of osteoporosis. Third, the follow-up time in the longitudinal analysis was five years on average, which might not be long enough to capture the cumulative effects of low-dose ICS use on BMD.

In conclusion, our study demonstrates that high-intensity use of ICS therapy slightly accelerates BMD loss at the hip in postmenopausal women with chronic respiratory diseases. However, this effect may not be clinically important as it would need to be sustained for over 40 years to produce one standard deviation reduction in BMD. It is important to balance concerns for the safety of ICS therapy with its effectiveness in reducing respiratory symptoms and improving quality of life in patients with COPD and asthma. This is especially the case for asthma, in which ICS therapy is the cornerstone of disease management^32,33^. The benefits of ICS for patients with COPD are likely more limited than have historically been reported, and recent data suggests that ICS may not be required in combination therapy to reduce the risk of exacerbations in certain patients^34^. Therefore, although our study does not support the discontinuation of long-term ICS therapy in post-menopausal women with chronic respiratory disease, its negative impacts on BMD in some patients warrants caution. Future studies should characterize the association between ICS use and the risk of fractures over a long follow-up period, as this is the final endpoint most relevant to the health of this population.

## ACKNOWLEDGEMENTS

The authors acknowledge the Manitoba Centre for Health Policy for use of data contained in the Manitoba Population Research Data Repository under HIPC Project Number 2011/2012-31). The results and conclusions are those of the authors and no official endorsement by the Manitoba Centre for Health Policy, Manitoba Health, or other data providers is intended or should be inferred. Data used in this study are from the Manitoba Population Research Data Repository housed at the Manitoba Centre for Health Policy, University of Manitoba and were derived from data provided by Manitoba Health. This article has been reviewed and approved by the members of the Manitoba Bone Density Program.

## Author Contributions

WDL, MS, and JMF formulated the study idea. WDL designed the study and performed all data analyses. JMF and MS contributed to the study design and interpretation of findings. WC and KJ and wrote the first draft of the manuscript (they are co-first authors). All authors critically commented on the manuscript and approved the final version. WDL is the guarantor of the manuscript.

